# The assembly and annotation of a new reference genome for the eastern mosquitofish (*Gambusia holbrooki*) reveal a new sex chromosome system in its invasive range

**DOI:** 10.64898/2026.11.18.767316

**Authors:** Alexandre Cormier, Kang Du, Florence Cornette, Antoine Fraimout, Eva Blondeau-Bidet, Arnaud Estoup, Manfred Schartl, Emilie Farcy, Charles Perrier, Céline Reisser

## Abstract

*Gambusia* species, especially eastern mosquitofish - *Gambusia holbrooki* - are of broad relevance to evolutionary biologists aiming at understanding genomic bases of adaptation during biological invasions and rapid response to selection to heterogeneous environments including pollution. Here we used HiFi and HiC to assemble a new, high quality, chromosome scale, genome sequence for *G. holbrooki* in its invasive range. Both haplotypes of the nuclear genome were assembled, comprising 657.64 Mb and 651.75 Mb respectively, in 24 chromosomes and had N50s of 28.97 Mb and 28.25 Mb. They had 94.87% and 94.29% BUSCO scores, illustrating a high completeness. Genome annotation yielded 25146 genes, which is comparable to other recent genome annotations of poecilid fish species. Repeat content was 25.69%, similar to what was previously reported for this species and for *G. affinis*. Synteny was highly conserved between the *G. holbrooki* assembly and the reference genome for *G. affinis* and *Xiphophorus maculatus* - shown here to be a genetically close species - with very few inter-chromosomal and little intra-chromosomal rearrangements. Mapping of resequenced males and females to the reference genome provided genomic evidence for a newly evolved homomorphic sex chromosome pair (W and Z, hence with female heterogamety) in *G. holbrooki*’s invaded range, which is unlike the XY system of its native range. Also, the new sex chromosomes are homologous to the ZW sex chromosome pair (chromosome 1) of the sister species *G. affinis*. This assembly provides a high quality reference genome for *Gambusia holbrooki*, in its invasive range in Europe, that is serving as a key resource for research on the genomics of invasion and of rapid adaptation to environmental change, pollution adaptation and sex determination evolution.

## Introduction & objectives

The mosquitofish species (*Gambusia* spp., particularly *G. affinis* and *G. holbrooki*) have emerged as model species in evolutionary ecology and ecotoxicology due to their high invasive potential, short generation time, and remarkable phenotypic and physiological plasticity, altogether resulting in a broad distribution (Pyke 2005, 2008). These features make mosquitofish especially suitable for investigating rapid adaptive responses to environmental stressors such as chemical pollutants, heavy metals, and endocrine disruptors (Díez-del- Molino *et al*. 2018), as well as the evolutionary processes underlying biological invasions, including dispersal, inbreeding, local adaptation and purge-accumulation of the genetic load. Access to a high-quality reference genome is essential to accurately identify genomic mechanisms implicated in evolutionary response during exposition to environmental variability, including pollution, and during invasion. Reference genomes have been generated for *Gambusia* species, among which chromosome level assemblies for *G. affinis* from China (Shao et al., 2020) and *G. holbrooki* from Italy (Shao *et al*. 2020, WELLCOME SANGER INSTITUTE 2025), which have already facilitated insights into the genomic basis of invasions and sex chromosome evolution (Vera, Díez-del-Molino, and García-Marín 2016, Kottler *et al*. 2020). New long-read sequencing technologies such as HiFi allow precise sequencing of long contigs, while HiC then allows precise chromosome scale scaffolding. In addition, the use of a local reference genome (assembled from individuals of the studied geographic region) mitigates reference bias in local studies, a well-known issue when relying on geographically/phylogenetically distant or generic references. Such bias can distort variant detection, diversity estimates, and signals of local adaptation, ultimately reducing the power of evolutionary population and functional genomic analyses, especially when geographic regions might have been colonised from different source populations (source of introduction). Hence, developing a local *Gambusia* reference genome for populations studied in a region of *G. holbrooki*’s invaded range, here the South of France, was a strategic investment to enhance both the resolution and accuracy of genomic inferences (Fraimout *et al*. 2026). Here we present the sequencing, assembly and annotation of a reference genome for *G. holbrooki* generated from a female specimen collected in Montpellier, Occitanie, France.

## Material and Methods

### DNA extraction, PacBio and Hi-C libraries preparation and sequencing

For the genome, we collected 13 females of *G. holbrooki* in PAU in April 2022 (Figure 1). One female (Figure 1A) was directly euthanised with an overdose of benzocaïne for PacBio HiFi library construction. The eyes, scales, gut and stomach were removed, and the remaining tissues were used for high molecular weight DNA extraction with the Macherey Nagel nucleobond High Molecular Weight kit, following manufacturer’s recommendations. HiFi library was constructed using SMRTbell® Template Prep kit 2.0. The library was loaded on one SMRTcell and sequenced on a Sequel III sequencer at the INRAE Genotoul GeT platform.

**Figure 1.**
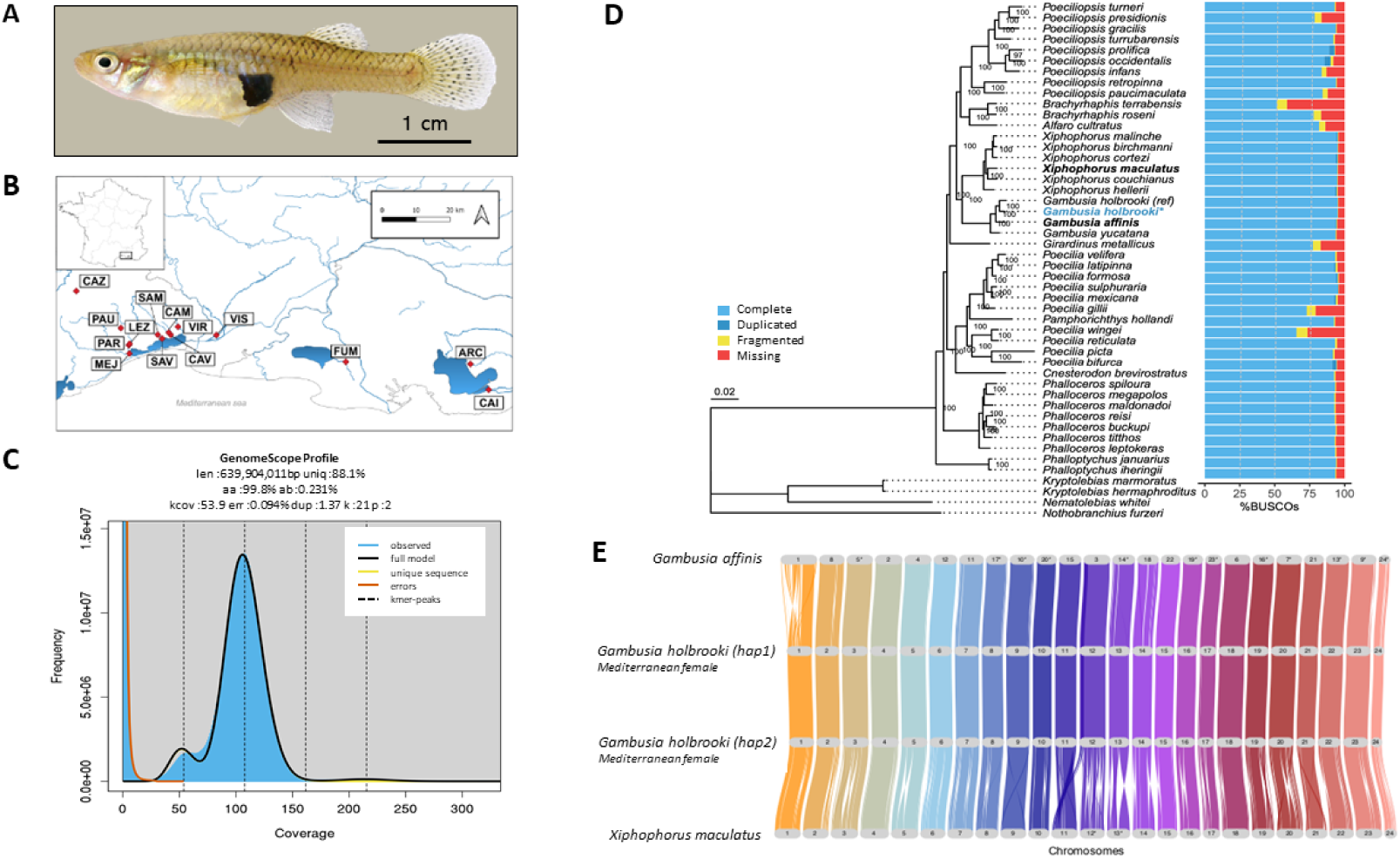
A) Female G. holbrooki specimens caught in the invasive range in France. B) Map of the sampled sites for genome assembly and sex chromosome identification (from Fraimout et al. 2026). C) GenomeScope profiling summary of the diploid genome, with the estimated genome size (len), and percentage of heterozygosity based on k-mer distributions. D) Maximum likelihood phylogenetic tree of genetic distance between *Gambusia holbrooki* (newly assembled genome indicated in blue with a star) and several other fish species, along with BUSCO completeness of the genomes. E) Synteny analysis indicating chromosomal rearrangements between the two newly assembled *Gambusia holbrooki* haplotypes and *G. affinis* and *Xiphophorus maculatus*.

The remaining 12 females were euthanised and their blood sampled from the dorsal vein after section of the caudal part, using a 10µL pipette with a sterile pipette tip coated with 10% EDTA solution. The extracted blood was pooled (14µL) and added to 2µL 10% EDTA to prevent coagulation, before being sent to the MGX platform for Hi-C library preparation. Libraries were constructed with the Arima-HiC kit from Arima Genomics and sequenced on the NovaSeq6000 using an SP flow cell and 150PE.

### Genome assembly and quality assessment

All scripts, pipeline used, and results for the assembly can be found at: https://bioinfo.gitlab-pages.ifremer.fr/bioanalysis/internal/gamboc/data/gamboc-female. We obtained 4558024 sequences from PacBio HiFi sequencing (Supplementary Table 1), with a N50=12397bp, and 278117542 PE sequences for Hi-C (Supplementary Table 1).

We used Cutadapt v4.1 (Martin 2011) to remove PacBio adapters (first: ATC TCT CTC AAC AAC AAC AAC GGA GGA GGA GGA AAA GAG AGA GAT, second: ATC TCT CTC TTT TCC TCC TCC TCC GTT GTT GTT GTT GAG AGA GAT) with a minimum overlap length of 35bp, and discarded reads with an error rate over 0.1.

We used HifiasmV0.19.7 (Cheng *et al*. 2021) to assemble a phased genome from HiFi and Hi-C reads, and used BandageNG v2022.08 (https://github.com/asl/BandageNG) for GFA visualisation. We assessed the quality of this first assembly using Biobox (Rudden *et al*. 2022) for metrics, Busco v5.5.0 (Simão *et al*. 2015) with the cyprinodontiformes ODB10 for completeness, and Merqury v1.3 (Rhie *et al*. 2020) for estimation of the completeness based on k-mer composition, using the exact method (Meryl). We purged duplications on both haplotypes using Purge_dups v1.2.6 (Guan *et al*. 2020), and performed QC analysis on this second assembly as described above. Finally, we used MitoHifi v3.4.1 (Uliano-Silva *et al*. 2023) to assemble the mitochondrial genome.

We aligned Hi-C reads to the purged assembly using BWA-MEM2 v2.2.1 (Vasimuddin *et al*. 2019) following the Arima mapping pipeline (https://github.com/ArimaGenomics/mapping_pipeline). We removed duplicate reads with Picard v2.21.1 (Broad 2019). We used YAHS v1.2a (Zhou, McCarthy, and Durbin 2023) to scaffold the assembly using Hi-C information. We performed manual curation of this first scaffolding using JBAT, following the author’s recommendations and tutorial (Dudchenko *et al*. 2018). We removed scaffolds presenting putative contamination (bacterial DNA) using MMseqs2 v14.7e284 (Steinegger and Söding 2017) and NCBI Foreign Contamination Screen v0.4.0 (O’Leary *et al*. 2023) from the AGP file and generated the decontaminated assembly using AGPtools v0.0.1 (Ricemeyer, Carroll, and Warren 2025). We finally organized the assembly using synteny with the *Xiphophorus maculatus* genome (GCF_002775205.1), using SeqKit v2.4.0 (Shen *et al*. 2016), minimap2 v2.24 (Li H 2021) and AGPtools v0.0.1.

### Genome structural and functional annotation

To characterize repetitive elements in the genome assembly, we generated a custom repeat library using RepeatModeler v2.0.5 (Flynn *et al*. 2020) with *-LTRStruct* and this repeat library was used to characterize the repeat contents of the assembly using RepeatMasker (within the RepeatModeler pipeline).

Genome assemblies were structurally annotated using an integrative framework adapted from established strategies (Powell *et al*. 2020, Du *et al*. 2024). This framework incorporated three complementary methodologies (comparative homology, transcriptome-guided annotation, and ab initio prediction) thereby ensuring a comprehensive and high-confidence gene catalog. Protein alignments and RNA-seq data provided evolutionary and transcriptional evidence, respectively, while gene structures were independently inferred de novo using AUGUSTUS v3.3.3 (Stanke *et al*. 2006). Subsequently, the different lines of evidence were synthesized into a unified annotation set.

For homology-driven annotation, a reference collection comprising 505310 protein sequences was downloaded. This dataset included vertebrate sequences from Swiss-Prot, the RefSeq “vertebrate_other” subset (restricted to accessions prefixed with “NP”), together with NCBI-annotated proteomes from human, zebrafish, platyfish, medaka, mummichog, turquoise killifish, guppy, and shortfin molly. Initial candidate loci were identified using GenblastA v1.0.4 (Birney, Clamp, and Durbin 2004), after which refined alignments were performed with GeneWise (included in GenblastA), while additional homology-based alignments were independently conducted using Exonerate v2.4.0 (https://github.com/nathanweeks/exonerate) and miniprot v0.18 (Birney, Clamp, and Durbin 2004, Li H 2023). When multiple models overlapped within a locus, the model with the highest alignment score was preferentially retained.

Transcriptomic evidence was integrated by mapping RNA-seq datasets derived from 36 experiments of two SRA bioprojects (Supplementary Table 2) using HISAT (Kim, Langmead, and Salzberg 2015). Transcript models were reconstructed with StringTie v2.2.3 (Pertea *et al*. 2015), while additional de novo assemblies were generated with Trinity v2.15.1 (Haas *et al*. 2013). These assemblies were subsequently aligned to the genome with Splign to infer exon–intron architecture with higher resolution (Kapustin *et al*. 2008).

Ab initio predictions were conducted using AUGUSTUS, which was subjected to a two-step training protocol. The initial training relied on BUSCO v5.8.2 in “--long” mode to generate a preliminary model, and the second phase utilized high-confidence gene models consistently recovered by GeneWise, Exonerate, StringTie, and Splign. Final AUGUSTUS predictions incorporated external hints derived from both homology- and transcript-based evidence. During evidence integration, gene loci were systematically reconciled across the three methodologies. In cases of conflicting predictions, priority was given to models most strongly supported by transcriptomic evidence. Exons with weak or ambiguous support were manually re-evaluated, and superior alternatives were incorporated where available. Predicted models lacking corroboration from either comparative or transcriptomic data were eliminated, resulting in the finalized, high-confidence gene annotation set.

The protein sequences of the final gene set were searched against the Swiss-Prot/UniProt database (https://www.uniprot.org/uniprotkb?query=reviewed:true) and the vertebrate subset of the RefSeq database (https://www.ncbi.nlm.nih.gov/refseq/) using BLAST (Camacho *et al*. 2009). Gene symbols and gene names were assigned based on the best hits, respectively. In addition, protein domains were annotated by searching against the Pfam database using InterProsSan (Paysan-Lafosse *et al*. 2023).

### Comparative genomics

Phylogenetic analyses were performed by comparing the BUSCO sequences from the *G. holbrooki* genome to those of 42 other Poeciliidae species, with 4 Aplocheiloidei genomes used as outgroups. External assemblies were retrieved from NCBI Datasets v18.7.0 (https://github.com/ncbi/datasets) using the corresponding taxonomic identifiers (Poeciliidae: 8079; Aplocheiloidei: 45443). Only reference genomes *(--reference*) were considered, and for Aplocheiloidei, assemblies were further restricted to those at the chromosome or complete level (*--assembly-level*; Supplementary Table 3 for all accession numbers). BUSCO analyses were conducted independently on each genome using *cyprinodontiformes* ODB10 dataset, which includes 15213 orthologous genes, in order to assess genomes completeness and then identify single-copy orthologs at the amino acid level. All single-copy BUSCO sequences from all genomes were concatenated into a single dataset, and only orthologs shared by all the genomes (48) were retained. In order to reduce the final alignment length, 150 ortholog groups were randomly selected among the 1578 shared between all the genomes. Multiple sequence alignments were generated for each of the 150 selected ortholog groups using MAFFT v7.520 (Katoh and Standley 2013). Poorly aligned or divergent regions were removed with Gblocks v0.91b (Talavera and Castresana 2007) with default parameters except for *-b5* set to *half*. The resulting curated alignments were concatenated into a supermatrix with ElConcatenero (https://odiogosilva.github.io/TriFusion/). The pipeline used can be found at: https://gitlab.ifremer.fr/bioinfo/workflows/pallor. Maximum likelihood phylogenetic inference was performed with IQ-TREE 2 v2.0.3 (Minh *et al*. 2020). Model selection was carried out automatically with ModelFinder limited to nuclear models (*-msub*), branch support was assessed using 1,000 ultrafast bootstrap replicates (*-B*) and 1000 replicates for SH approximate likelihood ratio test (*-alrt*) and the tree was rooted using Aplocheiloidei species. Tree rendering was done using ggtree v3.16.3 (Yu *et al*. 2017).

Global synteny relationship between our genome of *G. holbrooki* and the genomes of *G. affinis* and *Xiphophorus maculatus* were inspected with GENESPACE v1.2.3 (Lovell *et al*. 2022) using the gene positions derived from the GFF files, their corresponding orthogroups predicted with orthofinder v2.5.5 (Emms and Kelly 2018) and the syntenic blocks identified using MCScanX (Wang *et al*. 2012).

### Sex chromosomes identification

In order to identify the sex chromosomes, we resequenced the genome of a total of 66 individuals (33 males and 33 females) from 17 populations around the Gulf of Lion, France (Figure 1B). We additionally used published males and females RADseq sequences of the Florida *G. holbrooki* and *G. affinis* males and females, previously used in (Kottler *et al*. 2020) (SRA Bioproject accession number: PRJNA563257). Raw reads were processed with fastp v0.24.0 (Chen 2023), trimming remaining illumina adapters, and removing reads with an average quality under 20 and a length under 50bp. Trimmed reads were then mapped onto the newly assembled Haplotype 1 and Haplotype 2 of the mediterranean *G. holbrooki* with BWA-mem2 v2.3 (Vasimuddin *et al*. 2019) and default parameters. Mapped reads were trimmed with Sambamba v1.0.1 (Tarasov *et al*. 2015) to remove non properly paired reads, and reads mapping to multiple genomic locations. Remaining reads were used for variant calling using BCFTOOLS v1.23 (Danecek *et al*. 2021) and filter the resulting VCF with the filter functions to only keep bi-allelic variants with a minimum depth of 8, and less than 20% missing genotypes per locus. This pipeline was carried out separately for *G. holbrooki* from Florida, *G. affinis*, and for *G. holbrooki* from southern France which were sequenced using shotgun sequencing, resulting in three VCF files for each haplotype. We finally attempted to identify sex-linked genomic regions by calculating locus-based F_ST_ values between males and females using VCFTOOLS v0.1.16 (Danecek *et al*. 2011), and plotted the results using ggplot2 R package (Wickham 2016) to visualize locus-based F_ST_ values across all chromosomes, and identify putative sex determining regions (SDR).

### Chromosome 01 synteny and SDR microsynteny

To view sequence synteny across chromosome 01, we used the genome alignment module of Geneious V2025.2 (https://www.geneious.com) and the progressive Mauve algorithm (Darling *et al*. 2004) to align the mediterranean *G. holbrooki* haplotype 1 and 2 with *G. affinis* (Shao *et al*. 2020), *Xiphophorus maculatus* and *Poecilia reticulata* (Fraser *et al*. 2020). We used MCScanX (Wang *et al*. 2024) to identify syntenic gene blocks between both haplotypes across chromosome 01, and visualized the results using the online tool Synvisio (https://synvisio.github.io/#/dashboard/bn), with a single level analysis and the basic dashboard visualization.

We then focused on the microsynteny within the SDR region, and used minimap2 to align the 3.5Mb to 5Mb region of both haplotypes. Finally, we looked at TE content across chromosome 01 and in the SDR, for all TEs, and for young TEs (Kimura < 1, calculated through the RepeatModeler pipeline), and estimated LINE L1 integrity using TEsorter (Zhang *et al*. 2022) with the rexdb database.

## Results & discussion

### Genome assembly

Genomescope k-mer distribution analysis predicts a genome size of 640 Mbp and a 0.23% nucleotide heterozygosity (Figure 1C). Assembled haplotypes of the nuclear genome comprise 658.83 Mb and 651.19 Mb for haplotype 1 and haplotype 2 respectively (Supplementary files 1 and 2), with 24 chromosomes, matching the expectations from the cytogenetic karyotype of the Florida *G. holbrooki* (Müller *et al*. 2023) (Table 1). Haplotype 1 chromosomes have a N50 of 29.67 Mb, a N90 of 23.36 Mb, and a GC% of 39.06. Completeness of these haplotypes is high with 95.10% BUSCO for both haplotypes using the Cyprinodontiformes vODB10 database (Table 1, Figure 1D), and 97.3% and 96.6% using Merqury (Table 1). The final contact map shows a highly contiguous assembly (Supplementary Figure 1).

**Table 1.** Assembly statistics, BUSCO run with cyprinodontiformes_odb10, and Merqury statistics.

|  |  | **Haplotype 1** | **Haplotype 2** |
|---|---|---|---|
| Assembly statistics | Assembly length (bp) | 666,008,498 | 655,194,356 |
|  | Number of scaffolds | 24 chromosomes<br>+ 107 unplaced | 24 chromosomes<br>+ 46 unplaced |
|  | Scaffold chromosomes N50 (bp) | 29,674,054 | 28,543,450 |
|  | Number of contigs | 158 | 119 |
|  | Contig N50 (Mb) | 25,5 | 22,1 |
| BUSCO (n=15213) | Complete (C) | 14467 / 95.1% | 14306 / 94.0% |
|  | Complete single copy (S) | 14420 / 94.8% | 14262 / 93.7% |
|  | Complete duplicated (D) | 47 / 0.3% | 44 / 0.3% |
|  | Fragmented (F) | 54 / 0.4% | 59 / 0.4% |
|  | Missing (M) | 692 / 4.5% | 848 / 5.6% |
| Merqury | Completeness (%) | 97.32 | 96.58 |
|  | QV | 63.01 | 62.90 |

### Comparative genomics

Phylogenetic analysis based on the comparison of sequences of genes present in the Cyprinodontiformes vODB10 for 42 other poecilid species and 4 aplocheiloidei outgroup species revealed that *G. holbrooki* from southern France is very genetically very close to *G. holbrooki* from Padua, Italy (GCA_965282415.1 here labelled “ref”; Figure 1D) as well as *G. affinis* and *Xiphophorus* species, which is congruent with earlier findings based on mtDNA and partial nuclear gene sequences (Pollux *et al*. 2014) (Figure1D).

Chromosomal rearrangements in *G. holbrooki* compared to *G. affinis* and *X. maculatus* are rare and mainly intra-chromosomal, with the exception of chromosome 01 (the W sex chromosome of *G. affinis)*, showing large rearrangements and sequence loss (Figure 1E).

### Genome annotation

Repeat content identified by Repeat Modeler represented 25.69% and 24.67% of the genomic sequence of haplotype 1 and haplotype 2 respectively, among which 23.04% and 22.08% are made of TEs (details in Table 2). Gene annotation of haplotype 1 yielded 236,762 exons in 25,146 genes (Supplementary Table 4) and 258678 exons in 22513 genes for haplotype 2 (Supplementary Table 5), which is comparable to other recent genome annotations of poecilid fish species.

**Table 2.** Repeat annotations from RepeatModeler for haplotype 1 and haplotype 2.

|  | **Haplotype 1** |  |  | **Haplotype 2** |  |  |
|---|---|---|---|---|---|---|
|  | Number of elements | Length occupied | % of sequences | Number of elements | Length occupied | % of sequences |
| Retroelements | 117287 | 27056855bp | 4.06% | 115115 | 26238574bp | 4.03% |
| SINEs: | 10359 | 2057723bp | 0.31% | 10291 | 1979196bp | 0.30% |
| Penelope: | 0 | 0bp | 0.00% | 0 | 0bp | 0.00% |
| LINEs: | 82529 | 16708803bp | 2.51% | 81000 | 16271120bp | 2.50% |
| CRE/SLACS | 0 | 0bp | 0.00% | 0 | 0bp | 0.00% |
| L2/CR1/Rex | 58021 | 9892155bp | 1.49% | 57100 | 9705109bp | 1.49% |
| R1/LOA/Jockey | 2979 | 962883bp | 0.14% | 2960 | 966822bp | 0.15% |
| R2/R4/NeSL | 1167 | 244537bp | 0.04% | 1130 | 235088bp | 0.04% |
| RTE/Bov-B | 7601 | 2480100bp | 0.37% | 7361 | 2357673bp | 0.36% |
| L1/CIN4 | 8390 | 1642331bp | 0.25% | 8198 | 1622247bp | 0.25% |
| LTR elements: | 24399 | 8290329bp | 1.24% | 23824 | 7988258bp | 1.23% |
| BEL/Pao | 655 | 715266bp | 0.11% | 615 | 689649bp | 0.11% |
| Ty1/Copia | 48 | 30126bp | 0.00% | 45 | 32773bp | 0.01% |
| Gypsy/DIRS1 | 4150 | 3210972bp | 0.48% | 3932 | 3024308bp | 0.46% |
| Retroviral | 522 | 504669bp | 0.08% | 512 | 462301bp | 0.07% |
| DNA transposons | 381104 | 56469663bp | 8.48% | 377378 | 55750406bp | 8.56% |
| hobo-Activator | 120636 | 20203382bp | 3.03% | 119027 | 19916212bp | 3.06% |
| Tc1-IS630-Pogo | 206632 | 26524172bp | 3.98% | 205215 | 26301924bp | 4.04% |
| En-Spm | 0 | 0bp | 0.00% | 0 | 0bp | 0.00% |
| MULE-MuDR | 534 | 86899bp | 0.01% | 528 | 86211bp | 0.01% |
| PiggyBac | 1699 | 266763bp | 0.04% | 1692 | 264436bp | 0.04% |
| Tourist/Harbinger | 18393 | 3529499bp | 0.53% | 18136 | 3468112bp | 0.53% |
| Other (Mirage, P-element, Transib) | 0 | 0bp | 0.00% | 0 | 0bp | 0.00% |
| Rolling-circles | 10814 | 4142580bp | 0.62% | 10343 | 3968118bp | 0.61% |
| Unclassified: | 398368 | 69905908bp | 10.50% | 391951 | 61819842bp | 9.49% |
| Total interspersed repeats: |  | 153432426bp | 23.04% |  | 143808822bp | 22.08% |
| Small RNA: | 10115 | 2224054bp | 0.33% | 9896 | 1817290bp | 0.28% |
| Satellites: | 714 | 89880bp | 0.01% | 704 | 88098bp | 0.01% |
| Simple repeats: | 252606 | 11125666bp | 1.67% | 249466 | 10895661bp | 1.67% |
| Low complexity: | 35690 | 1654305bp | 0.25% | 35432 | 1631782bp | 0.25% |

### Sex chromosomes identification

Remapping of reads from males and females of *G. holbrooki* from Florida, *G. affinis*, and Mediterranean *G. holbrooki* from the present study revealed that the Mediterranean SDR is located on chromosome 01, which is syntenic to the *G. affinis* ZW chromosome pair, while *G. holbrooki* from Florida showed a clear signal on chromosome 16 (Figure 2A,B and C). The SDR signal (FST > 0.4) is restricted to a ∼59kb interval near the 4.45 Mb position on chromosome 01 (Figure 2D and E), and corresponds to a ZW sex determining system (Haplotype 1 being the Z chromosome), with heterozygous females and homozygous males, and a perfect segregation over 12kb (Figure 2E). This contrasts with *G. holbrooki* individuals from Florida, which exhibit a XY sex determining system (Kottler *et al*. 2020).

**Figure 2.**
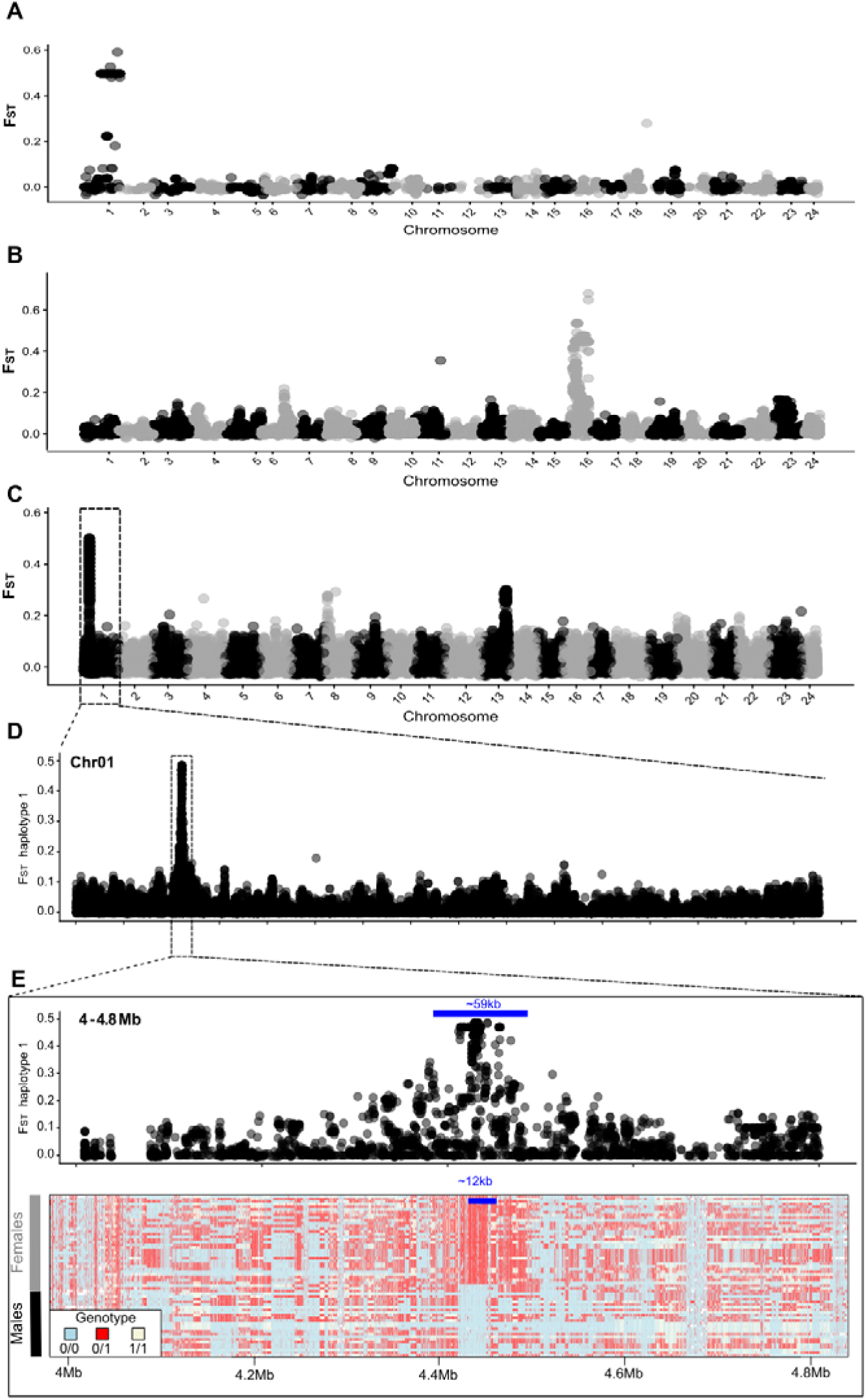
A) F_ST_ between males and females of *G. affinis*, B) F_ST_ between males and females of *G. holbrooki* from Florida, C) F_ST_ between males and females of *G. holbrooki* from the French mediterranean region, D) F_ST_ between males and females of *G. holbrooki* from the French mediterranean region along chromosome 1, E) F_ST_ between males and females of *G. holbrooki* from the French mediterranean region between 4Mb and 4.3Mb of the chromosome 1 (top panel), along with the genotype of the males and females (bottom panel).

### Chromosome 01 and SDR microsynteny

Although the Mediterranean ZW system mapped to the chromosome pair syntenic to the *G. affinis* ZW sex chromosomes, the W chromosome of the southern french population does not seem to be derived from *G. affinis*’s W chromosome, being more similar to *G. affinis*’s Z chromosome (Figure 3A). Global gene synteny of haplotype 1 and haplotype 2 is conserved, with only two small translocations and two small inversions on the second half of the chromosome (Figure 3B). Microsynteny of sequences showed that the SDR region is quite dynamic, with the presence of duplications and deletions in both haplotypes (Figure 3C). Microsynteny of genes showed that both haplotypes had events of duplications/deletions of a few genes, mostly uncharacterized genes, or genes related to transposable elements (Figure 4 top panel). The SDR region is preceded in both haplotypes by a TE dense region (Figure 4), annotated as young LINEs L1-Tx1 (Figure 4 bottom panel), which do not occur at that density in the rest of the genome (Supplementary Figure 2). Haplotype 1 had more young LINEs than haplotype 2, and while more than 70% of the young LINEs of haplotype 1 exhibited all the ORFs of active TEs, those of haplotype 2 were truncated, with only 40% retaining full integrity (Table 3).

**Figure 3.**
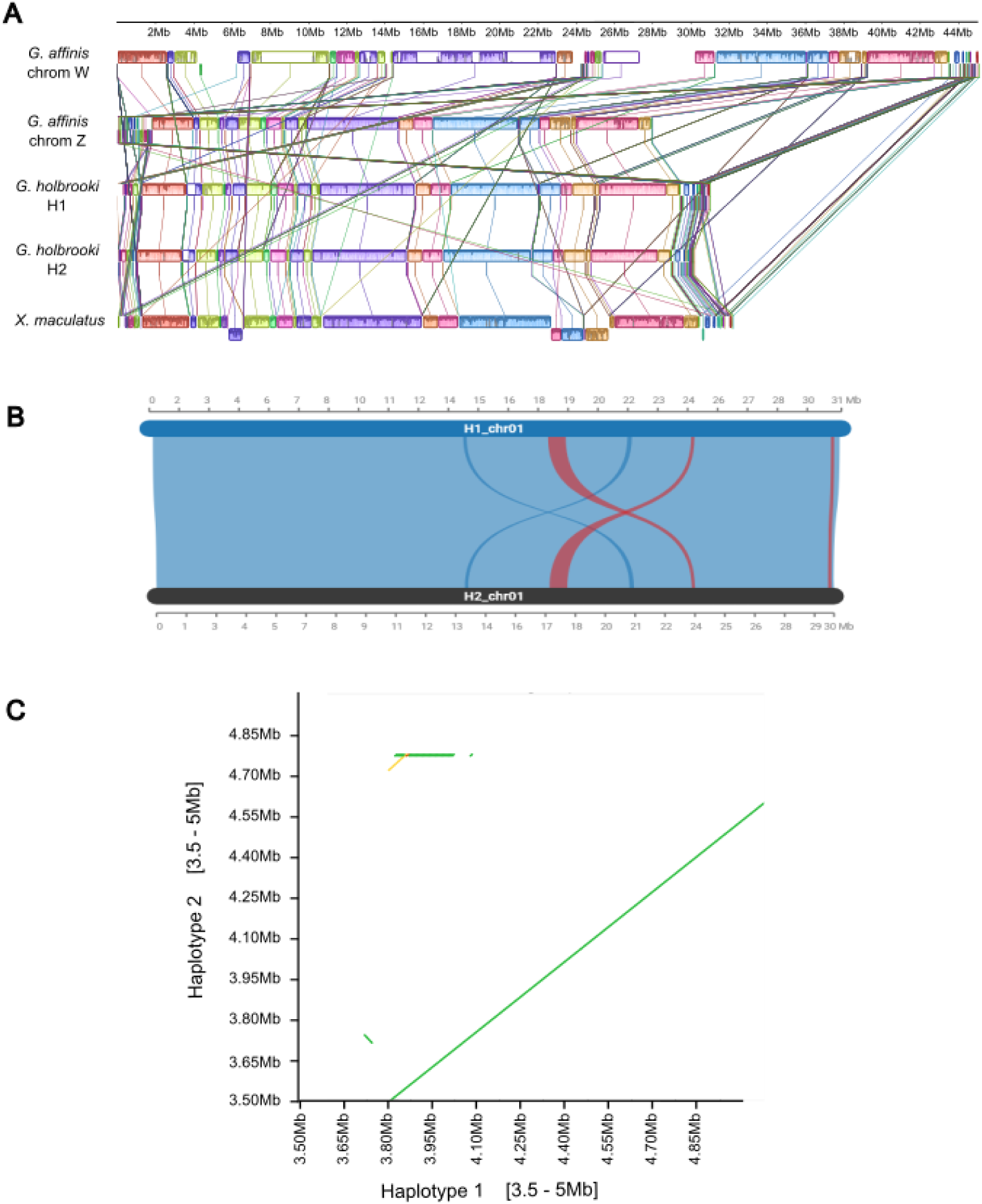
(A) Sequence macrosynteny of Chromosome 01, using haplotype 1 and haplotype 2 of the mediterranean *G. holbrooki*, chromosome 01 of *Xiphophorus maculatus*, and the W and Z of *G. affinis*. (B) Gene synteny between haplotype 1 and haplotype 2 of the mediterranean *G. holbrooki* chromosome 01. Dark blue strings represent translocation and red strings represent inversions (with or without translocation); (C) Sequence microsynteny in the SDR region of haplotype 1 and haplotype 2 of the mediterranean *G. holbrooki*.

**Figure 4.**
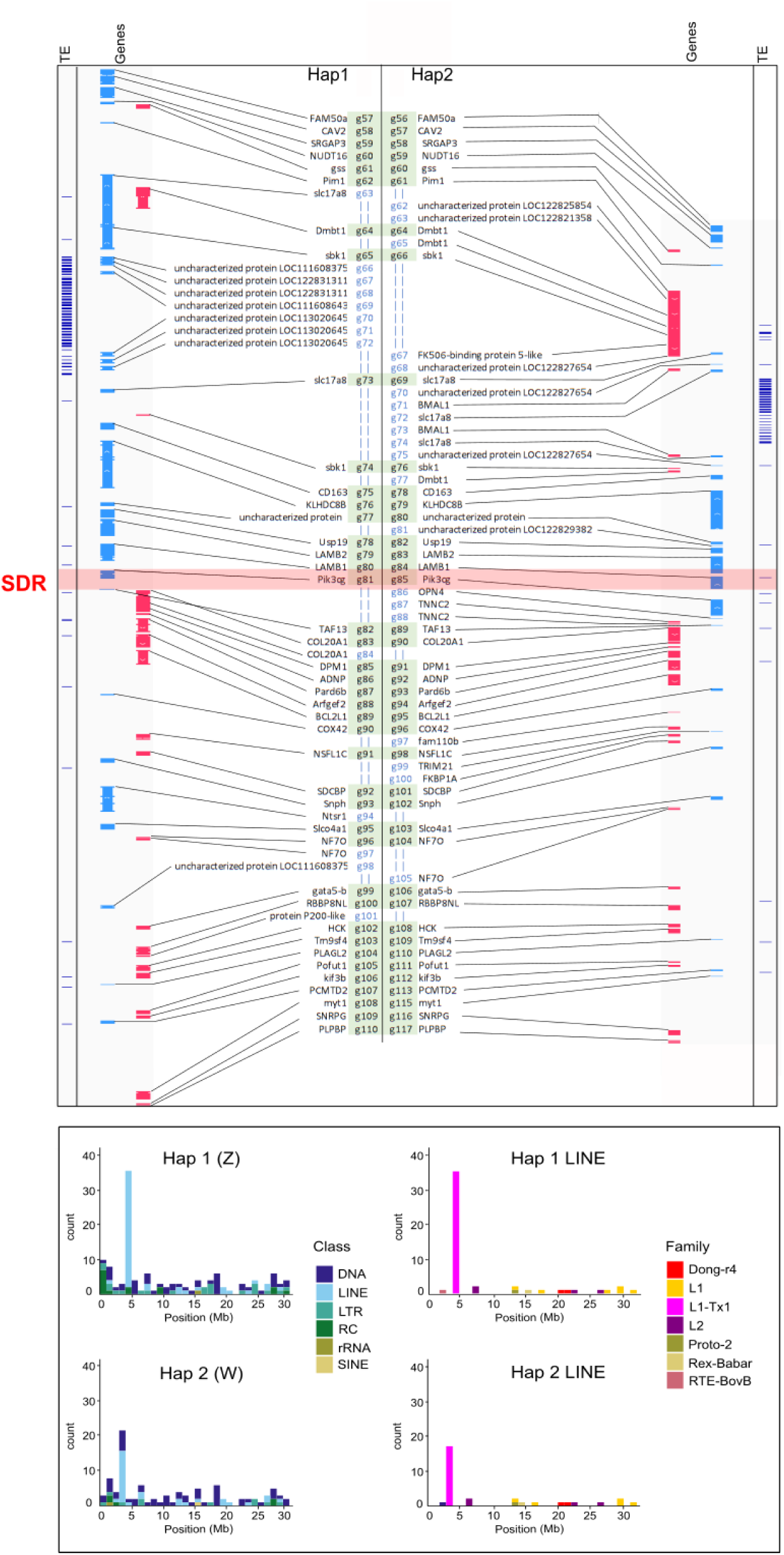
Gene synteny around the SDR region of the mediterranean *G. holbrooki* (top), and distribution of TE classes and LINE families along chromosome 01 in haplotype 1 and haplotype 2. The sex determining region is indicated by a red rectangle.

**Table 3.** Percentage of the LINE L1-Tx1 near the SDR region on haplotype 1 and haplotype 2 that shows complete integrity (conservation of the three typical motifs: for endonuclease, LINE retrotransposase, and LINE ORF).

| **Domains** | **Haplotype 1** | **Haplotype 2** |
|---|---|---|
| ENDO \| RT \| LINE | 71,11 | 40,00 |
| ENDO \| LINE | 2,22 | 0,00 |
| RT \| LINE | 2,22 | 5,71 |
| No_Domain | 24,44 | 54,29 |

Finally, the SDR region showing perfect patterns of heterozygosity (all males homozygous, all females heterozygous) contains the phosphatidylinositol-4,5-bisphosphate 3-kinase catalytic subunit gamma gene (PIK3CG; Figure 4A). Phosphoinositide 3-kinases (PI3Ks) do phosphorylate inositol lipids and are involved in the immune response (Figure 4 top panel). Two neighbour genes LAMB1 and LAMB2, might be of interest. Indeed, LAMB1 is known to be differentially expressed between male and female gonads in rainbow trout (Depiereux *et al*. 2015). LAMB2 is also known to be involved in gonadal differentiation and reproduction in fish, and is linked to the Wnt signalling pathway (Li *et al*. 2026).

## Conclusion and perspectives

This reference genome of *Gambusia holbrooki* provides a critical resource for investigating the genomic bases and mechanisms underlying rapid adaptation to novel environments in invasive populations of this species. In southern France, *G. holbrooki* thrives in a wide range of extreme habitats, including highly polluted and occasionally anoxic waters contaminated with heavy metals, lipophilic organic compounds, and pharmaceuticals, as well as hypersaline environments where salinity can reach up to 56 g·L ¹ (Martin *et al*. 2025, Fraimout *et al*. 2026). The availability of this genomic resource will also enable detailed comparative analyses of the recently identified sex-determining system in southern French populations with that of populations from the northern United States, particularly in the Potomac region, which is believed to represent the source of the European introductions. Together, these resources open new opportunities to explore the genomic mechanisms underlying environmental tolerance, invasion success, and the evolution of sex-determination systems in this globally invasive species.

## Data availability

Scripts used for the genome assembly are available from https://bioinfo.gitlab-pages.ifremer.fr/bioanalysis/internal/gamboc/data/gamboc-female/. The raw sequencing data as well as haplotype 1 and haplotype 2 assemblies and gene annotations are available in the European Nucleotide Archive (ENA) under BioProject accession number PRJEB129762 (https://www.ebi.ac.uk/ena/browser/view/PRJEB129762).

## Ethics and sampling permits

### Funding

The project was funded by the Occitanie Regional Council’s program « Key 543 challenge BiodivOc » (GambOc project) to E. Farcy. MGX acknowledges financial support from France 544 Génomique National infrastructure, funded as part of the “Investissement d’Avenir” program 545 managed by Agence Nationale pour la Recherche (contract ANR-10-INBS-09).

### Authors contribution

AC, AF, AE, MS, CP, EF and CMOR designed the experiment. FC and EBB performed DNA extractions. AC performed genome assembly and phylogenetic analysis. KD performed genome structural and functional annotation. CMOR performed comparative genomic and population genomics analyses. All authors contributed to the redaction of the manuscript.

## Supplementary materials

**Supplementary File 1**: Assembly of Haplotype 1of the mediterranean *G. holbrooki*.

**Supplementary File 2**: Assembly of Haplotype2 of the mediterranean *G. holbrooki*.

**Supplementary Table 1.** Statistics on HiFi and Hi-C raw data.

|  | **Nb seq** | **Sum Length (bp)** | **Min Length (bp)** | **Average Length (bp)** | **Max Length (bp)** | **N50** | **Q30(%)** | **GC(%)** |
|---|---|---|---|---|---|---|---|---|
| **PacBio HiFi** | 4558024 | 70416139563 | 93 | 15448.80 | 57518 | 12397 | 92.97 | 38.83 |
| **Hi-C R1** | 278117542 | 41717631300 | 150 | 150 | 150 | 150 | 91.95 | 40.86 |
| **Hi-C R2** | 278117542 | 41717631300 | 150 | 150 | 150 | 150 | 90.85 | 40.91 |

**Supplementary Table 2.**
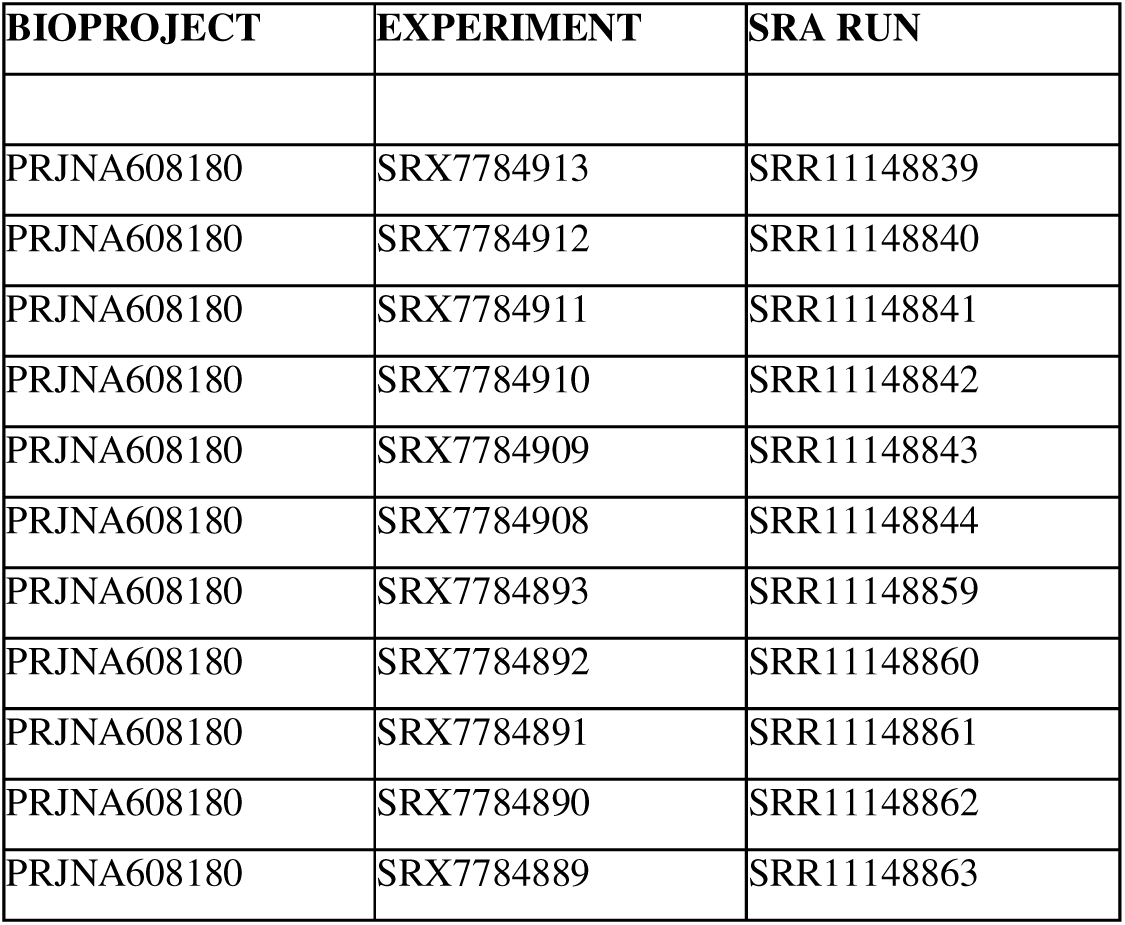
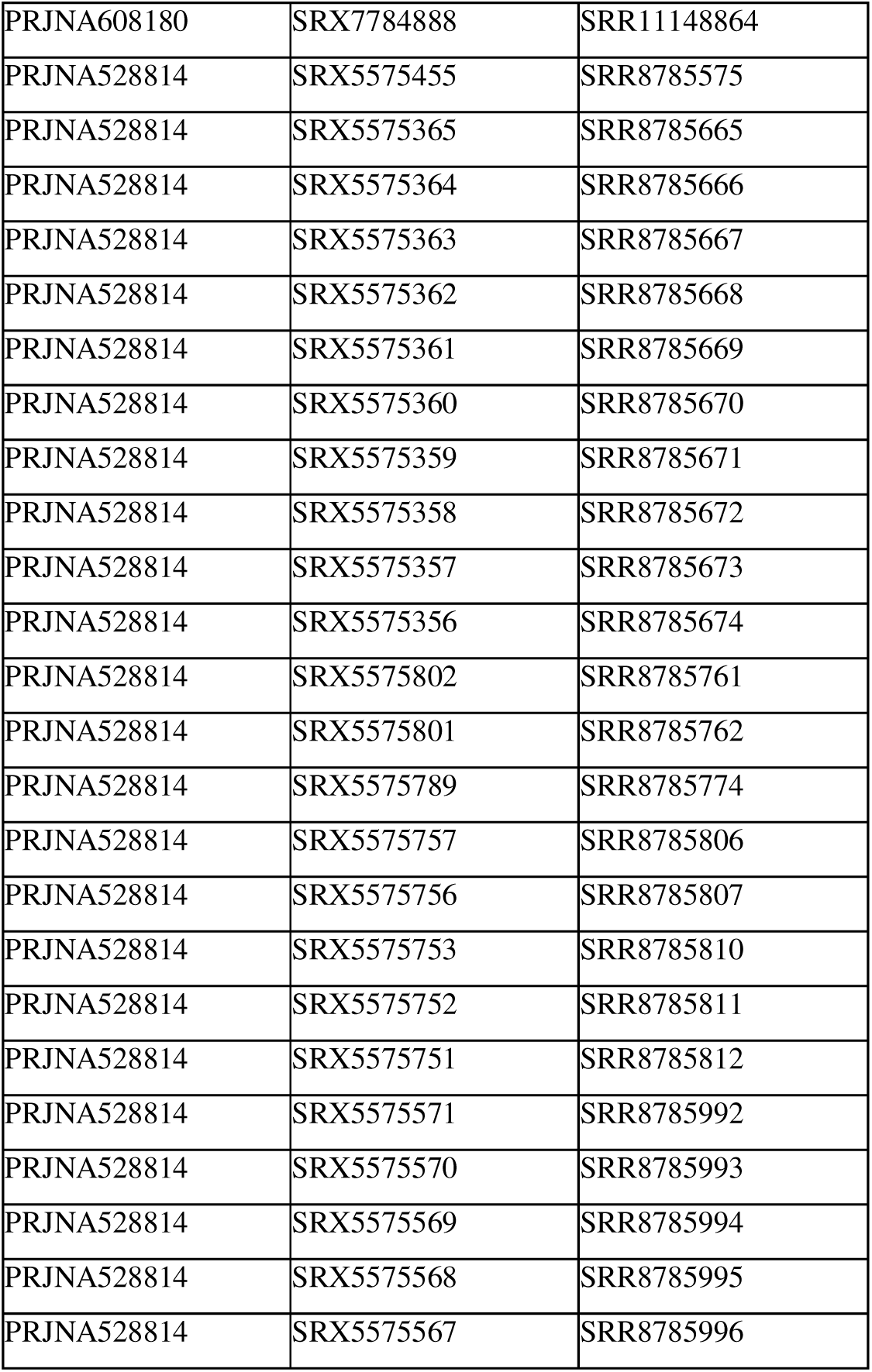
Accession numbers of SRA reads used for functional annotation.

**Supplementary Table 3.** Accession numbers for phylogenetic analysis and comparative genomics.

| NCBI Accession | Species |
|---|---|
| GCA_903066825.1 | *Alfaro_cultratus* |
| GCA_903066815.1 | *Brachyrhaphis_roseni* |
| GCA_903067065.1 | *Brachyrhaphis_terrabensis* |
| GCA_046269305.1 | *Cnesterodon_brevirostratus* |
| GCF_019740435.1 | *Gambusia_affinis* |
| GCA_965282415.1 | *Gambusia_holbrooki (ref)* |
| GCA_035584275.1 | *Gambusia_yucatana* |
| GCA_903067095.1 | *Girardinus_metallicus* |
| GCA_007896545.1 | *Kryptolebias_hermaphroditus* |
| GCF_001649575.2 | *Kryptolebias_marmoratus* |
| GCF_014905685.2 | *Nematolebias_whitei* |
| GCF_043380555.1 | *Nothobranchius_furzeri* |
| GCA_046118195.1 | *Pamphorichthys_hollandi* |
| GCA_046118275.1 | *Phalloceros_buckupi* |
| GCA_046118015.1 | *Phalloceros_leptokeras* |
| GCA_046118075.1 | *Phalloceros_maldonadoi* |
| GCA_046118095.1 | *Phalloceros_megapolos* |
| GCA_046118115.1 | *Phalloceros_reisi* |
| GCA_046118235.1 | *Phalloceros_spiloura* |
| GCA_046118035.1 | *Phalloceros_titthos* |
| GCA_046118055.1 | *Phalloptychus_iheringii* |
| GCA_902982915.1 | *Phalloptychus_januarius* |
| GCA_052054765.1 | *Poecilia_bifurca* |
| GCF_000485575.1 | *Poecilia_formosa* |
| GCA_903068115.1 | *Poecilia_gillii* |
| GCF_001443285.1 | *Poecilia_latipinna* |
| GCF_001443325.1 | *Poecilia_mexicana* |
| GCA_033032245.1 | *Poecilia_picta* |
| GCF_000633615.1 | *Poecilia_reticulata* |
| GCA_051168745.1 | *Poecilia_sulphuraria* |
| GCA_036873795.1 | *Poecilia_velifera* |
| GCA_903068105.1 | *Poecilia_wingei* |
| GCA_911649635.1 | *Poeciliopsis_gracilis* |
| GCA_903067105.1 | *Poeciliopsis_infans* |
| GCA_010883035.1 | *Poeciliopsis_occidentalis* |
| GCA_903067055.1 | *Poeciliopsis_paucimaculata* |
| GCA_903068095.1 | *Poeciliopsis_presidionis* |
| GCF_027474105.1 | *Poeciliopsis_prolifica* |
| GCA_010277075.1 | *Poeciliopsis_retropinna* |
| GCA_922788865.1 | *Poeciliopsis_turneri* |
| GCA_010277015.1 | *Poeciliopsis_turrubarensis* |
| GCA_036418095.1 | *Xiphophorus_birchmanni* |
| GCA_041380905.1 | *Xiphophorus_cortezi* |
| GCF_001444195.1 | *Xiphophorus_couchianus* |
| GCF_003331165.1 | *Xiphophorus_hellerii* |
| GCF_002775205.1 | *Xiphophorus_maculatus* |
| GCA_041429825.1 | *Xiphophorus_malinche* |

**Supplementary Figure 1.**
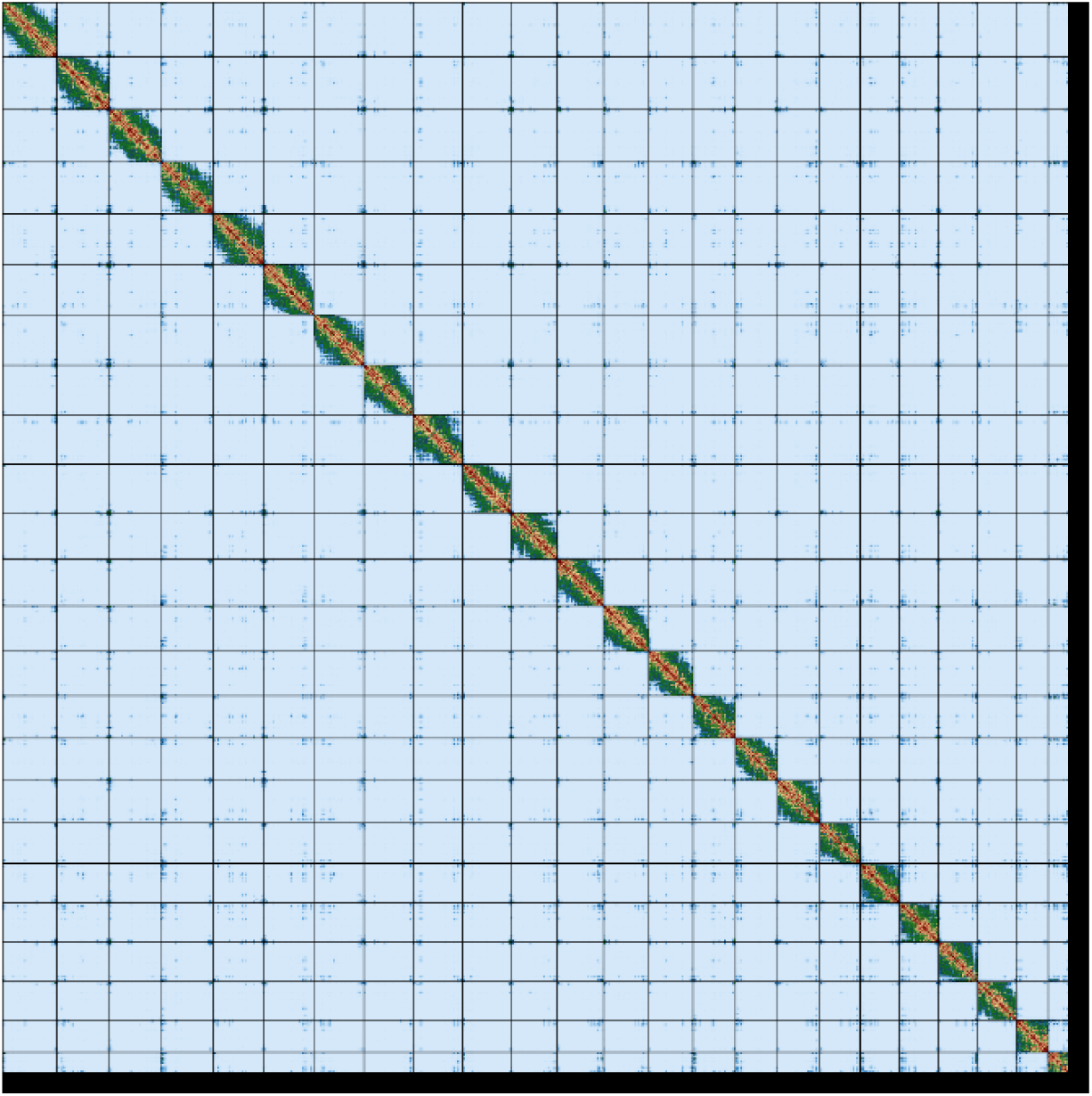
Contact map of Hi-C assemblyof the G. holbrooki genome.

**Supplementary Figure 2.**
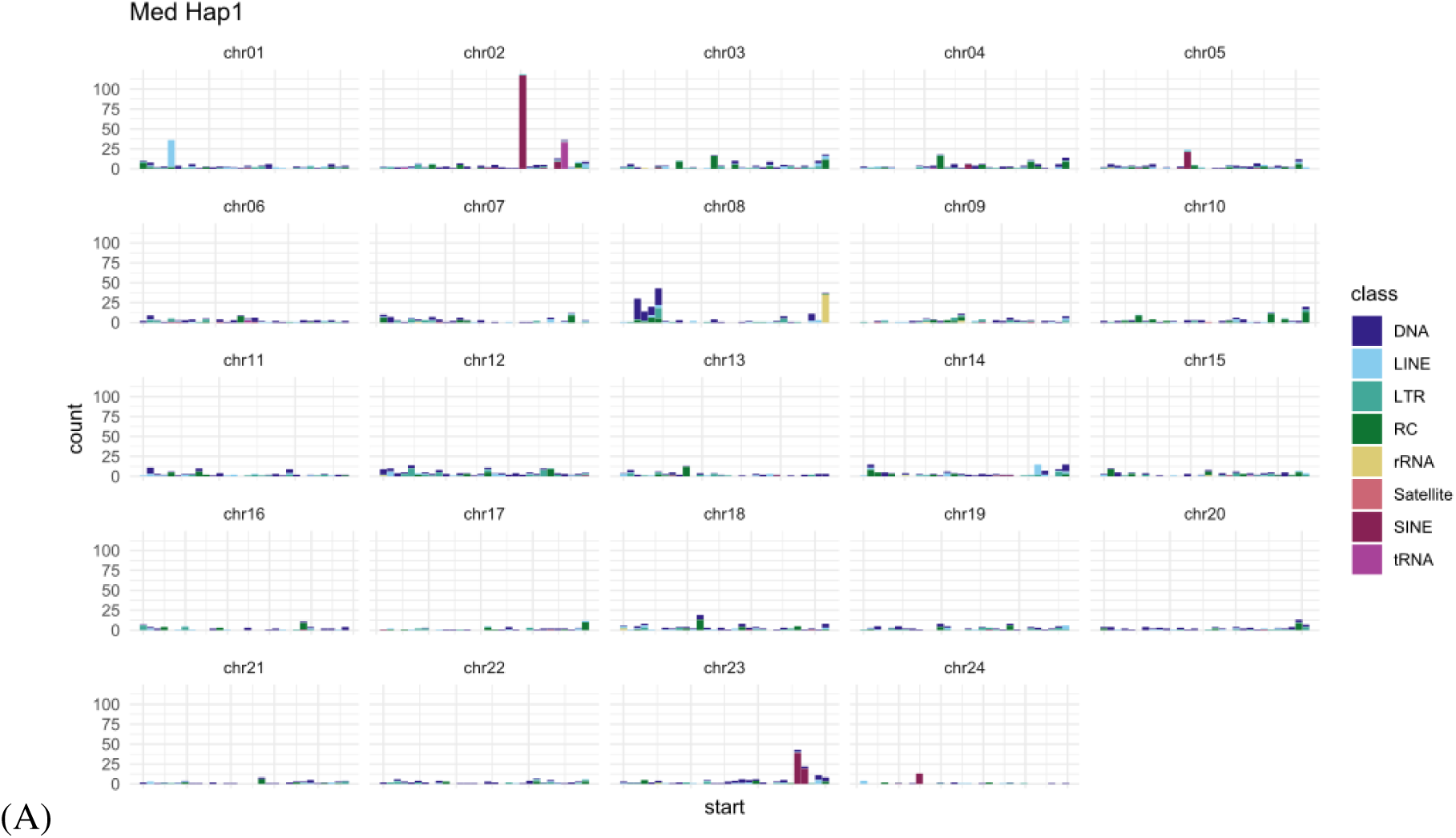
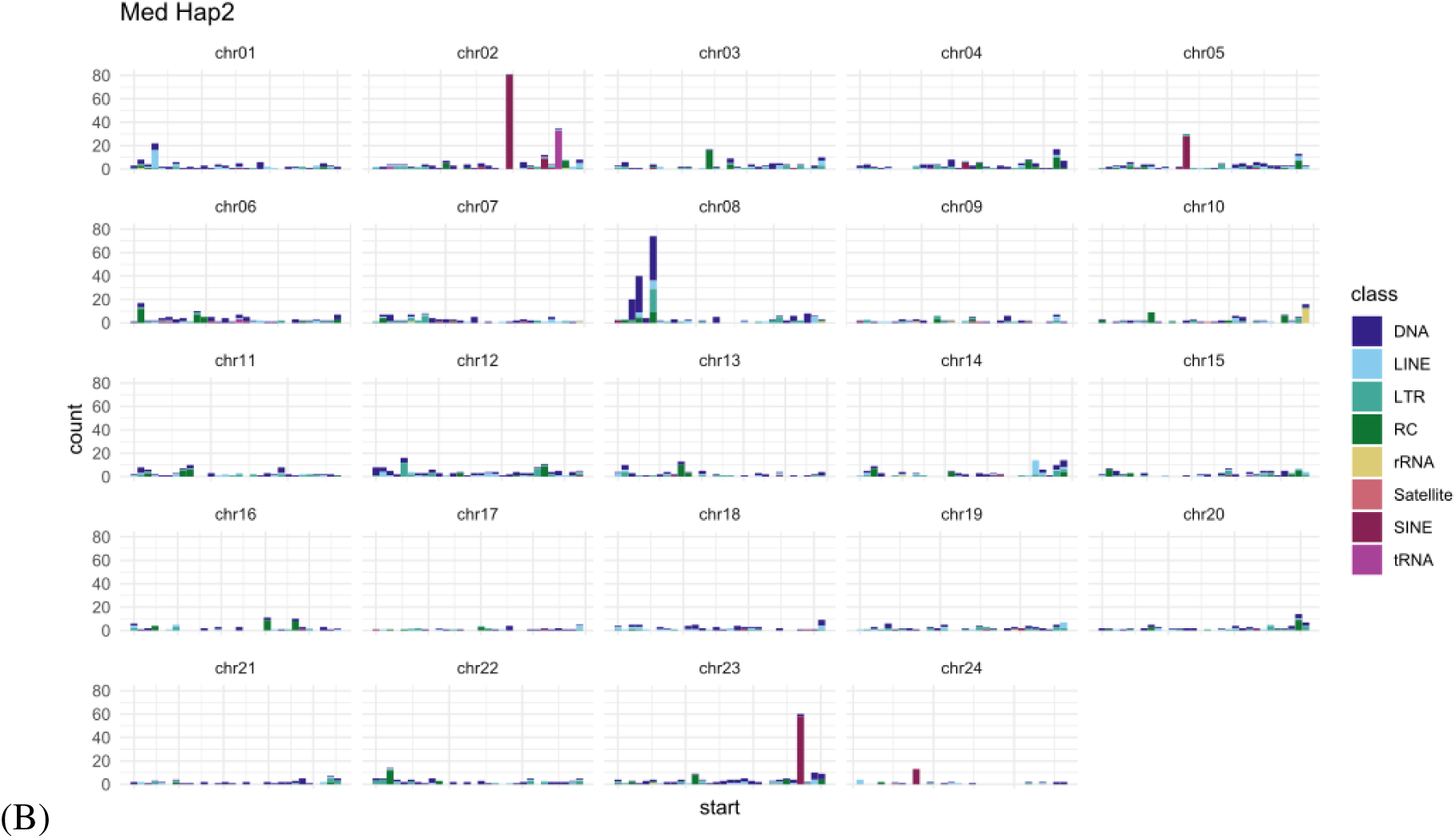
Young >TE content (Kimura < 1) for all chromosomes in (A) haplotype1, (B) haplotype 2.

